# DNA accessibility is not the primary determinant of chromatin-mediated gene regulation

**DOI:** 10.1101/639971

**Authors:** Răzvan V. Chereji, Peter R. Eriksson, Josefina Ocampo, David J. Clark

## Abstract

DNA accessibility is thought to be of major importance in regulating gene expression. We test this hypothesis using a restriction enzyme as a probe of chromatin structure and as a proxy for transcription factors. We measured the digestion rate and the fraction of accessible DNA at all genomic *Alu*I sites in budding yeast and mouse liver nuclei. Hepatocyte DNA is more accessible than yeast DNA, consistent with longer linkers between nucleosomes, and indicating that nucleosome spacing is a major determinant of accessibility. DNA accessibility varies from cell to cell, such that essentially no sites are accessible or inaccessible in every cell. *Alu*I sites in inactive mouse promoters are accessible in some cells, implying that transcription factors could bind without activating the gene. Euchromatin and heterochromatin have very similar accessibilities, suggesting that transcription factors can penetrate heterochromatin. Thus, DNA accessibility is not likely to be the primary determinant of gene regulation.

## INTRODUCTION

Genomic DNA is packaged into chromatin, which is composed of regularly spaced nucleosomes. Human and mouse cells contain relatively open euchromatin (similar to yeast chromatin) and extremely compact heterochromatin (Becker et al. 2016; Allshire and Madhani 2018). Most genes located in heterochromatin are completely repressed (Becker et al. 2016). Controlling the accessibility of DNA to transcription factors is thought to be of major importance in regulating gene activation, primarily through precise positioning of a nucleosome over regulatory elements such as promoters and enhancers, blocking access to transcription factors (Fig. 1A). Activation is thought to occur when an ATP-dependent chromatin remodeler removes the blocking nucleosome, allowing transcription factors to bind, although precisely how remodelers are targeted to regulatory elements is still unclear (Voss and Hager 2014; Venkatesh and Workman 2015).

**Figure 1.**
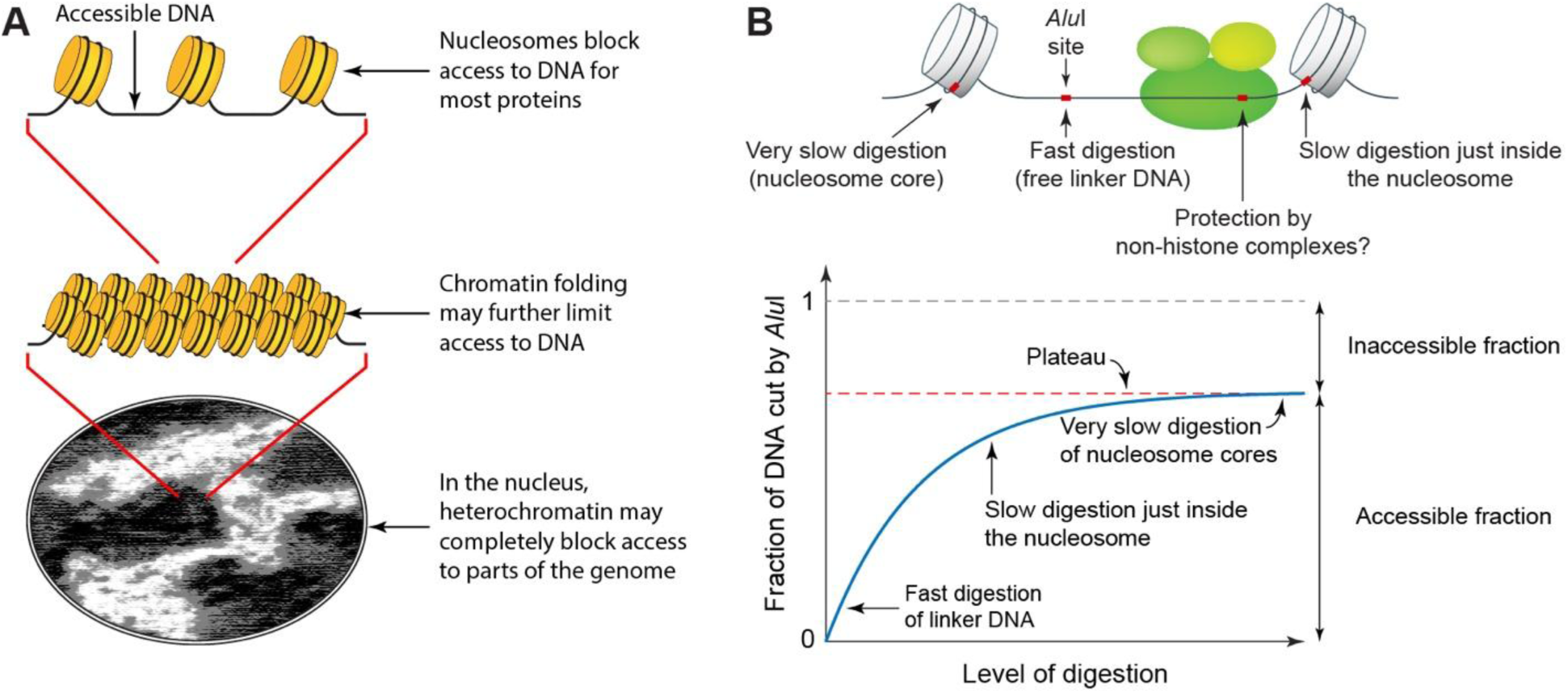
A fully quantitative assay for DNA accessibility in chromatin. (A) DNA accessibility may depend on the degree of chromatin compaction. (B) Principle of the restriction enzyme protection assay.

DNA accessibility may also be modulated at higher levels of chromatin structure (Fig. 1A): linker histone-dependent condensation of the chromatin filament may limit access to linker DNA between nucleosomes. Furthermore, large-scale compaction may occlude transcription factors from heterochromatin domains, perhaps involving liquid droplet phase separation (Larson et al. 2017). If DNA accessibility is the primary determinant of gene regulation, then it is crucial for repression that critical regulatory elements are blocked in *all* cells in a population. Otherwise, there would be inappropriate gene activation in some cells.

The accessibility model described above is appealing, but it has not yet been tested using a fully quantitative genome-wide assay. MNase-seq data are difficult to quantify because micrococcal nuclease destroys the DNA as digestion proceeds. Furthermore, nucleosomes are digested at different rates depending on the sequences they contain, resulting in apparently different relative occupancies as digestion proceeds (Chereji et al. 2017; Schwartz et al. 2018). MNase-seq data are typically normalized to the genomic average and relative nucleosome occupancies are estimated, although they are still subject to the caveat above. Three other valuable methods, DNase-seq (John et al. 2011), ATAC-seq (which uses a transposase) (Schep et al. 2015) and RED-seq/NA-Seq (which use restriction enzymes) (Gargiulo et al. 2009; Chen et al. 2014) report the relative accessibilities of open regions in chromatin. Small DNA fragments excised from accessible DNA sequences (typically regulatory elements) are isolated and sequenced. However, the rest of the genome is excluded from the analysis because it is still present as very long DNA molecules. Consequently, the signal and relative amounts of each accessible element depend on the extent of digestion, allowing only relative measurements. Since these methods sequence only the small fraction of accessible DNA fragments, they are only semi-quantitative.

We have adapted the restriction enzyme protection assay to measure accessibility (Linxweiler and Hörz 1984; Fascher et al. 1990; Jack and Eggert 1990; Archer et al. 1991; Verdin et al. 1993; Wallrath and Elgin 1995; Shen et al. 2001). This assay measures both the absolute accessibility of the DNA and the rate at which accessible sites are cut. It has been used *in vitro* to monitor nucleosome reconstitution (Zheng and Hayes 2003), to detect nucleosome shifts (Studitsky et al. 1994) and to measure the activities of chromatin remodeling enzymes (Tsukiyama and Wu 1995). It depends on the fact that restriction enzymes are essentially unable to cut their cognate sites within a nucleosome (Linxweiler and Hörz 1984; Polach and Widom 1995). Restriction enzymes cut nucleosomal DNA 10^2^ - 10^5^ times slower than linker DNA, with faster rates for DNA just inside the nucleosome, as it is more likely to dissociate transiently from the histone octamer (Polach and Widom 1995; Chereji and Morozov 2014). These large rate differences result in a plateau in the digestion, corresponding to the fraction of accessible DNA (Fig. 1B; Supplemental Text).

Here, we use the restriction enzyme protection assay to measure the accessibility of a large number of specific sites throughout the genome in nuclei from budding yeast and mouse liver. We show that essentially no sites are blocked in every cell or accessible in every cell, demonstrating a high degree of cell-to-cell heterogeneity, and inconsistent with nucleosome block models. In addition, we find that DNA in budding yeast chromatin is less accessible than DNA in mouse hepatocyte chromatin, which is explained by the shorter nucleosome spacing in yeast and suggests that nucleosome spacing is the primary determinant of DNA accessibility. Mouse hepatocyte euchromatin and heterochromatin have very similar absolute DNA accessibilities and are penetrated at very similar rates, indicating that heterochromatin does not prevent restriction enzymes from accessing their sites and presumably would not exclude transcription factors either.

## RESULTS

### A fully quantitative measure of DNA accessibility: qDA-seq

We used the restriction enzyme *Alu*I to measure both the absolute DNA accessibility (i.e. the fraction of the DNA that is accessible to *Alu*I) and the initial rate at which these accessible sites are cut. This simple method, which we term “<u>q</u>uantitative <u>D</u>NA <u>a</u>ccessibility” assay (qDA-seq), involves treating nuclei with a restriction enzyme at different concentrations, sonicating the DNA into small fragments, followed by paired-end sequencing. The sonication step is necessary because the *Alu*I digest contains many long DNA fragments derived from protected chromatin, which are not suitable for Illumina sequencing.

*Alu*I cuts the sequence AG|CT to yield blunt ends. The yeast genome has ∼40,000 *Alu*I sites; the mouse genome has ∼12.6 million sites. After sequencing, we calculate the fraction of DNA molecules cut at each *Alu*I site as a function of *Alu*I concentration up to ∼50 nM. The accessible fraction is measured by counting the number of DNA molecules with an end corresponding to a specific genomic *Alu*I site as a fraction of all DNA molecules containing the same site. A direct comparison of accessible fractions in different cell types is possible; critically, no normalization is necessary. We note that although most of the protection observed is likely to be due to nucleosomes, the nucleosome may not be the only complex that is resistant to restriction enzymes. Such complexes would have to be stable enough to protect an *Alu*I site during the entire incubation (20 minutes).

### DNA accessibility in yeast varies from cell to cell

To avoid potential complications due to increased accessibility of replicating DNA, we arrested haploid yeast cells in the G1 phase of the cell cycle using α-factor. Nuclei were digested with increasing concentrations of *Alu*I and the expected plateau was observed at essentially all *Alu*I sites (see below). It is important to note that only one copy of a unique genomic sequence is present in each cell, because the cells are haploid. Therefore, the plateau value indicates the fraction of cells in which a particular unique *Alu*I site is accessible. Each site is accessible in some cells and inaccessible in the other cells.

We present the *ARG1* gene as an example (Fig. 2A). Digestion at an *Alu*I site (site 2; Fig. 2A) just inside the −1 nucleosome reaches a plateau at ∼45% cut, indicating that this site is accessible in ∼45% of the cells and inaccessible in the remaining ∼55% of cells. A neighbouring *Alu*I site (site 3) located close to the upstream border of the nucleosome-depleted region (NDR) at the *ARG1* promoter is more accessible, at ∼50%. In contrast, all three *Alu*I sites in the *ARG1* coding region (sites 4, 5 and 6; located within the +2, +3 and +7 nucleosomes respectively) have lower accessibilities (∼15-20%), suggesting that nucleosome occupancy is higher on the coding region (∼75-80%), consistent with MNase-seq data (Fig. 2A), and indicating that these sites are accessible in only 1 in 4 or 5 cells. The *Alu*I site in the *YOL057W* promoter downstream of *ARG1* (site 8) is much more accessible, but the digestion still reaches a plateau at ∼60%, indicating that this site is protected in ∼40% of cells, even though the MNase-seq data show that it is located within a deep NDR, predicting a nucleosome occupancy close to 0 (Fig. 2A). Instead, we attribute this protection to non-histone proteins stably bound at the *YOL057W* promoter in ∼40% of cells (Chereji et al. 2017). Importantly, a plateau is reached at all *Alu*I sites, indicating that each site is accessible in some cells and protected in the remaining cells.

**Figure 2.**
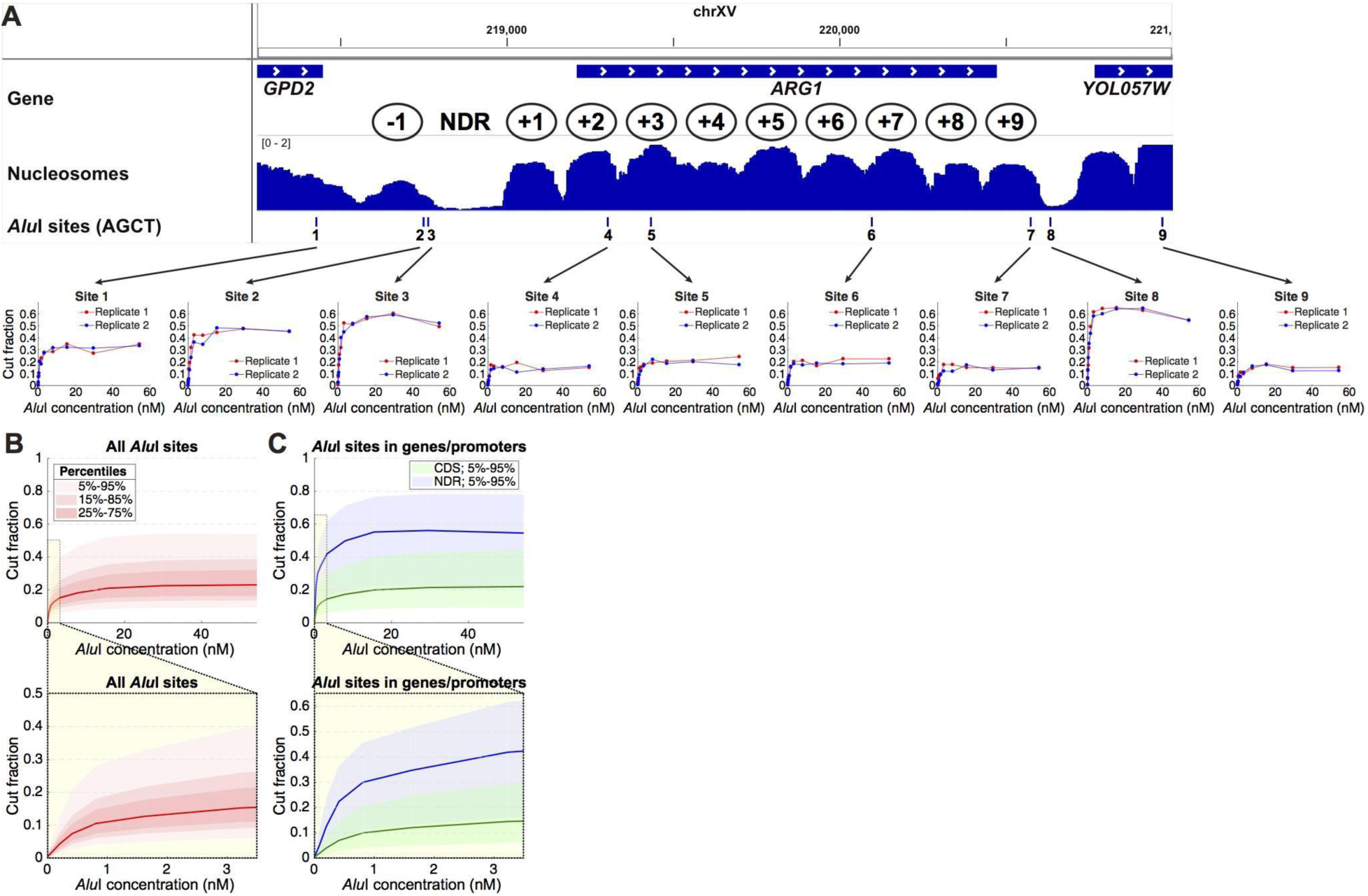
DNA accessibility in yeast varies from cell to cell. (A) *Alu*I accessibility of the *ARG1* gene in arrested haploid yeast cells. Upper panel: Nucleosome occupancy (MNase-seq data (Ocampo et al. 2016)) in wild type cells normalized to the genomic average (= 1). Ovals indicate approximate nucleosome positions. Lower panel: *Alu*I digestion at each of nine sites (data for two independent experiments are shown). The plateau value is a measure of the fraction of cells in which the *Alu*I site is accessible. Each site is accessible in some cells and inaccessible in the rest. (B) Digestion kinetics for all ∼40,000 *Alu*I sites as a function of [*Alu*I] (11 digestion points) for haploid cells arrested with α-factor. Red line: median level of digestion. Pink shading indicates data ranges: the lightest pink area includes 90% of the *Alu*I sites (*i.e*. the 5%-95% data range, which excludes the 5% of *Alu*I sites that are the least cut and the 5% of sites that are the most cut). Lower panel: Initial stages of digestion. (C) Kinetics for *Alu*I sites in gene bodies (between start and stop codons) and promoter NDRs defined using the positions of the +1 and −1 nucleosomes (Chereji et al. 2018). Blue line: median level of digestion in NDRs; green line: median level of digestion in gene bodies.

For genome-wide analysis of the data, we superimposed the plots for all ∼40,000 *Alu*I sites (Fig. 2B). A plateau is reached at a median accessibility of ∼22%, indicating that the median *Alu*I site is inaccessible in ∼78% of cells. The data range shows that 90% of *Alu*I sites are cut in only ∼9% - 55% of cells (Fig. 2B). This observation indicates that yeast cells are very heterogeneous in DNA accessibility. To gain more insight, we divided the *Alu*I sites into gene body sites and promoter sites (Fig. 2C). The median accessibility in gene bodies is very similar to that for all sites (∼20%), because gene bodies account for a very large fraction of the yeast genome. If we make the simple assumption that a nucleosome protects 147 bp of every 165 bp (the average nucleosome spacing in yeast (Thomas and Furber 1976)), the predicted protection is 89% (147/165), which is higher than observed (∼80%), suggesting that there may be some digestion just inside the nucleosome as observed *in vitro* (Polach and Widom 1995), or that there may be occasional gaps in the nucleosomal arrays. In the former case, a value of 80% is consistent with a protected inner nucleosome core of 132 bp (80% of 165 bp), suggesting that the outer ∼7 bp on both sides of the nucleosome are vulnerable to *Alu*I.

The median accessibility of promoters, defined by their NDRs, is ∼53% (range: 90% of sites cut in 22% - 78% of cells), which is much higher than in gene bodies, and consistent with nucleosome depletion. However, digestion within the NDR still reaches a plateau, indicating the presence of stable complexes protecting the NDR in about half of the cells, presumably corresponding to non-histone barrier complexes, as discussed above. Estimation of *Alu*I digestion rates at *accessible* sites in promoter NDRs and genes, assuming first order kinetics (see Supplemental Text), indicates that NDR sites are digested only ∼1.3 times faster than linker DNA sites in gene bodies (Fig. 2B, C; Supplemental Fig. S1).

### Imperfect nucleosome positioning can account for cell-to-cell heterogeneity in DNA accessibility

We plotted the mean *Alu*I accessibility for all ∼5,000 yeast genes as a function of distance from the first (+1) nucleosome on the gene, which typically covers the transcription start site (TSS) in yeast (Mavrich et al. 2008) (Fig. 3A). The extent of digestion as a function of *Alu*I concentration is shown. In the absence of *Alu*I, there is a background of ∼1% cut, corresponding to random fragmentation of the DNA at an *Alu*I site by sonication. Digestion increases with increasing *Alu*I concentration up to ∼15 nM, beyond which there is no more digestion. There is a strong peak at the NDR, with a maximum mean value of ∼55% cut. Promoters are more accessible than gene bodies, but a resistant complex is present in about half of the cells.

**Figure 3.**
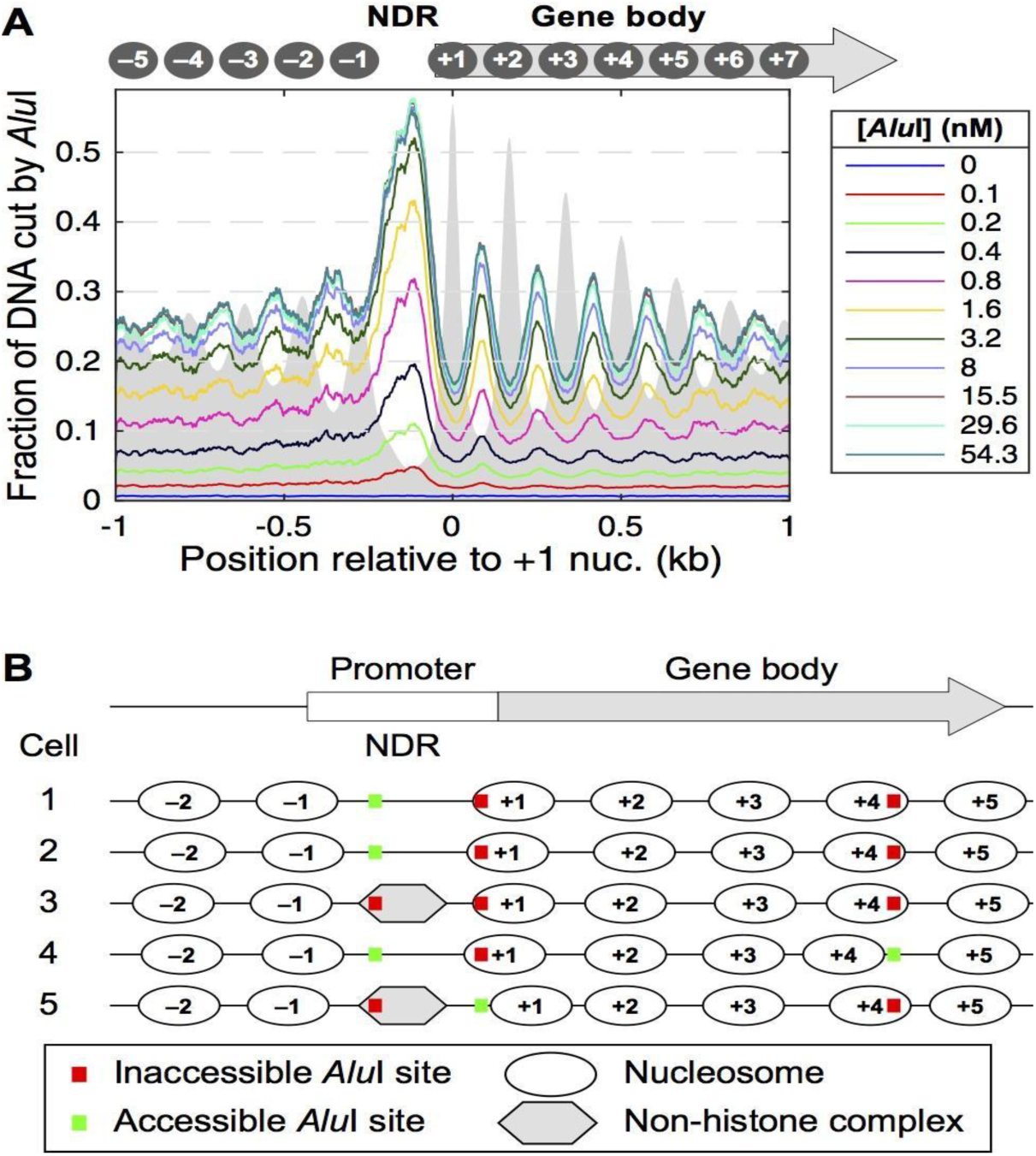
Genomic analysis of *Alu*I accessibility reveals imperfect nucleosome phasing in yeast. (A) Mean accessibility as a function of distance from the center of the +1 nucleosome (defined by Chereji et al. 2018) on all ∼5,000 yeast genes. (B) Heterogeneous nucleosome positioning model to explain the *Alu*I accessibility data. On a typical gene, nucleosomes are positioned slightly differently in each cell such that a particular *Alu*I site is inside a nucleosome in one cell and in a linker in another cell. The cartoon shows the nucleosome positions on a gene in five different cells. An *Alu*I site in the coding region is in the linker (accessible) in only one cell out of five (20% accessibility), whereas an *Alu*I site in the promoter NDR is accessible in three out of five cells (60% accessibility). The observed average values are ∼25% in the coding region and ∼55% in the NDR (see A).

In gene bodies, an oscillatory pattern is observed around a mean value of ∼25% accessibility, anti-correlated with phased nucleosomes observed in MNase-seq data, such that the *Alu*I peaks coincide with linkers and the *Alu*I troughs coincide with nucleosomes. The amplitude of this oscillation provides a quantitative estimate of the degree of phasing. Perfectly phased nucleosomal arrays (*i.e*., each nucleosome occupies an identical position in every cell) predict 100% cutting at *Alu*I sites in linkers (*i.e*., cut in all cells) and 0% cutting at nucleosomal sites (*i.e*., blocked in all cells). In fact, the oscillations are relatively weak: the average probability of cutting an *Alu*I site located at the +1 nucleosome position is ∼15%, compared to ∼35% for *Alu*I sites in linkers. Thus, the +1 nucleosome is shifted or absent in ∼15% of cells. These data can be explained by a model in which regularly spaced nucleosomes are positioned slightly differently in different cells, such that an *Alu*I site is protected in ∼80% of cells and located in an accessible linker in ∼20% of cells (Fig. 3B). Similarly, promoters are blocked by a stable complex in about half of the cells.

### Heavy transcription correlates with increased DNA accessibility of yeast gene bodies

We treated exponentially growing yeast cells with 3-aminotriazole (3AT), which induces the amino acid starvation response mediated by the Gcn4 transcription factor (Hinnebusch and Natarajan 2002). We have shown previously that 3AT induces heavy transcription of *ARG1, HIS4* and a few other genes, resulting in chromatin disruption and loss of nucleosome occupancy over the coding region and flanking regions (Cole et al. 2014). As expected, the *Alu*I accessibility of the *ARG1* and *HIS4* gene bodies increases after 3AT treatment, whereas the accessibility of *Alu*I sites in *GAL1*, which is not induced by 3AT, is unaffected (Supplemental Fig. S2). We also note that growing cells and α-factor arrested cells have similar DNA accessibilities at the global level, suggesting that replication does not have a strong effect on global accessibility (Supplemental Fig. S2; compare with Fig. 3A).

### The mouse hepatocyte genome is more accessible than the yeast genome

We performed the same experiment using mouse liver nuclei. *Alu*I digestion resulted in a plateau at a median accessibility of ∼34% (Fig. 4A). Thus, the mouse hepatocyte genome is more accessible than the yeast genome (∼22%; Fig. 2B). Although this observation seems counter-intuitive, given that the yeast genome is very active and lacks heterochromatin, it is consistent with the longer average nucleosome spacing in hepatocytes (∼195 bp; van Holde 1989) relative to yeast (∼165 bp). More insight is obtained by examining the chromatin structure in the vicinity of the average mouse promoter after alignment of all ∼25,000 genes on the TSS (Fig. 4B). *Alu*I digestion in the promoter NDR just upstream of the TSS reaches a plateau at ∼45% accessibility, which is higher than in the flanking regions (∼32% accessible/ ∼68% protected). The protection of genic DNA is consistent with a protected inner nucleosome core of 133 bp (68% of 195 bp), which is essentially the same as that observed for yeast genes (132 bp). Weak nucleosome phasing is apparent downstream.

**Figure 4.**
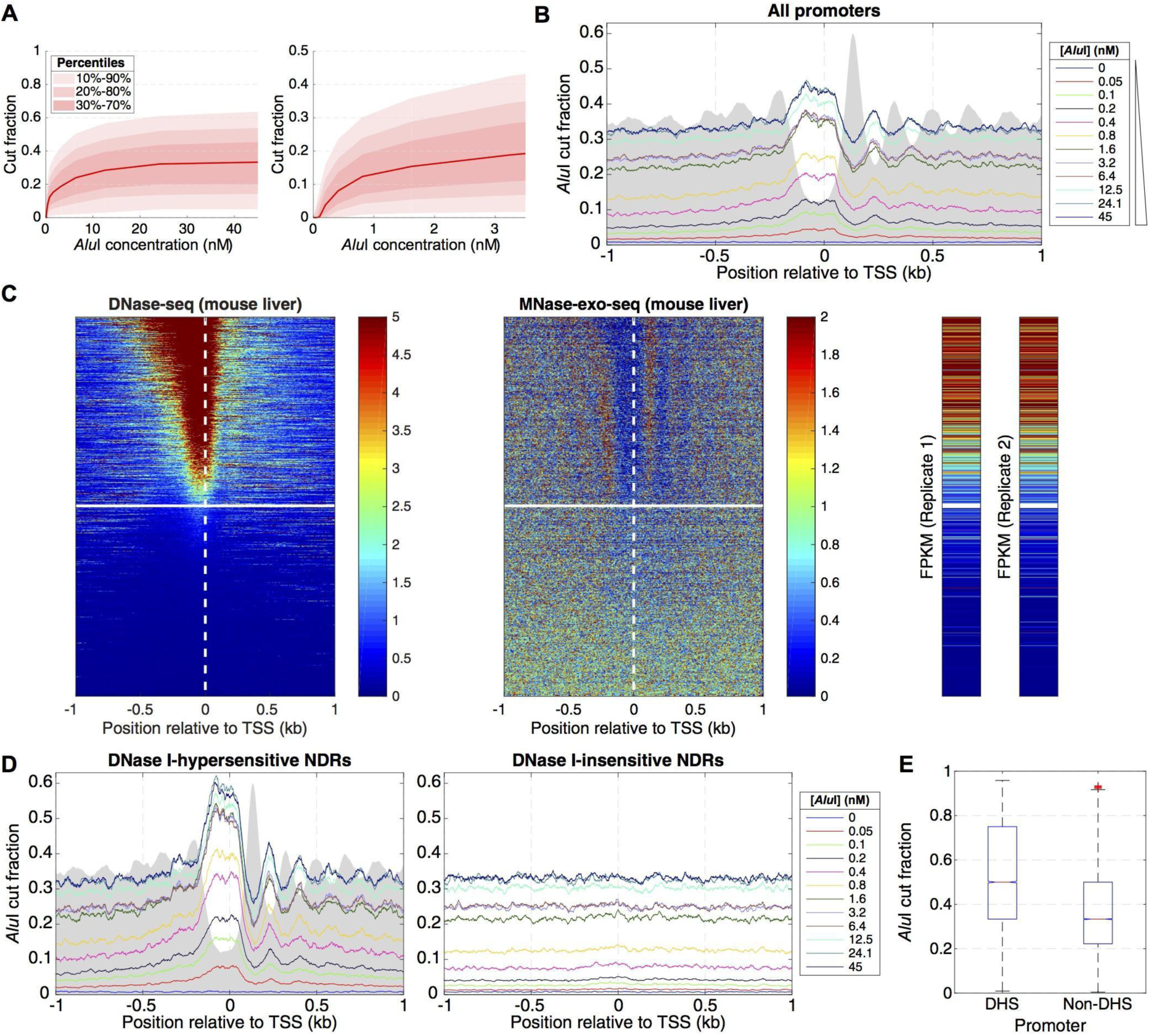
Inactive gene promoters are accessible to *Alu*I in some mouse liver cells. (A) *Alu*I digestion of mouse hepatocyte nuclei (12 digestion points) Left panel: all data. Right panel: initial digestion. The data range shows that 80% of *Alu*I sites (with at least 5 reads) are cut in 5% - 60% of cells. (B) Average *Alu*I accessibility plotted as a function of distance from the TSS on all ∼25,000 mouse genes. Grey area: MNase-Exo-seq data (nucleosome dyads (Cole et al. 2016)) on an arbitrary scale. (C) Heat map analysis of all ∼25,000 mouse genes sorted according to the DNase I hypersensitivity of their promoters in mouse hepatocytes (data from ENCODE) and aligned on the TSS: left panel: DNase I cut density; middle: nucleosome dyad positions (Cole et al. 2016); right: RNA-seq data (two biological replicates from ENCODE). The white line divides hypersensitive and insensitive promoters (defined in Supplemental Fig. S3). (D) Average *Alu*I accessibility plotted as a function of distance from the TSS for DNase I-hypersensitive and DNase I-insensitive promoters defined in C. (E) Distribution of *Alu*I cut fractions corresponding to sites located in promoters (region [TSS - 185 bp; TSS + 85 bp]), separated by DNase I hypersensitivity. The difference in DNA accessibility is highly significant (two-sided Kolmogorov-Smirnov test (test statistic = 0.316) and two-sided t-test (test statistic = 96.288); p-values < 10^-4^; confidence interval for the difference in population means, at significance level 0.05: [0.158; 0.164]; effect size: Cohen’s d = 0.755).

### Inactive mouse gene promoters are accessible in some cells

We sorted the genes according to the DNase I hypersensitivity of their promoters in mouse hepatocytes. This analysis revealed two classes of promoter: hypersensitive and insensitive (Fig. 4C; Supplemental Fig. S3A) (Chereji and Clark 2018). After sorting using the same gene order, nucleosome positioning (MNase-Exo-seq) data (Cole et al. 2016) and hepatocyte gene expression data show that genes with hypersensitive promoters are mostly active, with an NDR and phased nucleosomes, whereas genes with DNase I-insensitive promoters are inactive, lack phasing and have no NDR (Fig. 4C; Supplemental Fig. S3).

Analysis of the *Alu*I data indicates that active genes show better phasing and higher NDR accessibility (∼58%) than all genes (Fig. 4D; *cf*. Fig. 4B). On the other hand, DNA accessibility on both sides of the NDR is unchanged (∼32%; Fig. 4D). In contrast, DNase I-insensitive genes are uniformly accessible (∼32%), including promoters, with no evidence for an NDR or nucleosome phasing, consistent with the nucleosome positioning data (Fig. 4C). Although inactive promoters have a lower mean accessibility (∼32%) than active promoters (∼58%), they are at least partly accessible in ∼1 in 3 haploid genomes (i.e. on at least one allele in half of these diploid cells). The surprising accessibility of *Alu*I sites in inactive promoters probably reflects the lack of nucleosome phasing, such that regularly spaced, but unphased nucleosomal arrays result in protection of promoter *Alu*I sites in cells where they are nucleosomal and exposure in the other cells where they are in linker DNA (*cf*. Fig. 3B).

### Euchromatin and heterochromatin have very similar DNA accessibilities

We compared the accessibilities of euchromatin and heterochromatin using a 15-state epigenetic model for mouse hepatocyte chromatin derived from histone modification patterns and ChIP-seq data for Pol II and CTCF (Bogu et al. 2015) (Fig. 5). Surprisingly, the median absolute *Alu*I-accessible fraction is similar for all 15 chromatin states (the plateau values range from ∼29% to ∼36%), indicating that all states are accessible, including all heterochromatin states and Polycomb-repressed regions (Fig. 5A). Active promoters (states 5 and 7) and strong enhancers (states 6 and 8) were defined in the model of Bogu et al. (2015) primarily by the presence of the H3-K4me1, H3-K4me3, H3-K27ac histone marks and Pol II, whereas insulators were defined primarily by CTCF binding (state 15). All three of these regulatory elements are more accessible to *Alu*I than the euchromatin and heterochromatin states, because they are short and dominated by an NDR, which has a higher average accessibility than the flanking chromatin (Fig. 4D, left panel). (Note that the accessibility of active promoters (∼36%; Fig. 5A) averages lower than at promoter NDRs (58%; Fig. 4D), because the epigenetic state model includes both the NDR and its modified flanking nucleosomes.) Most importantly, the curves for the euchromatin states (1 and 2), defined by the H3-K36me3 mark (Bogu et al. 2015), track with those for the heterochromatin states, defined by the H3-K27me3 mark (Polycomb-repressed; state 11) or by the absence of active marks (states 12 - 14), indicating that the differences between them are negligible (Fig. 5A).

**Figure 5.**
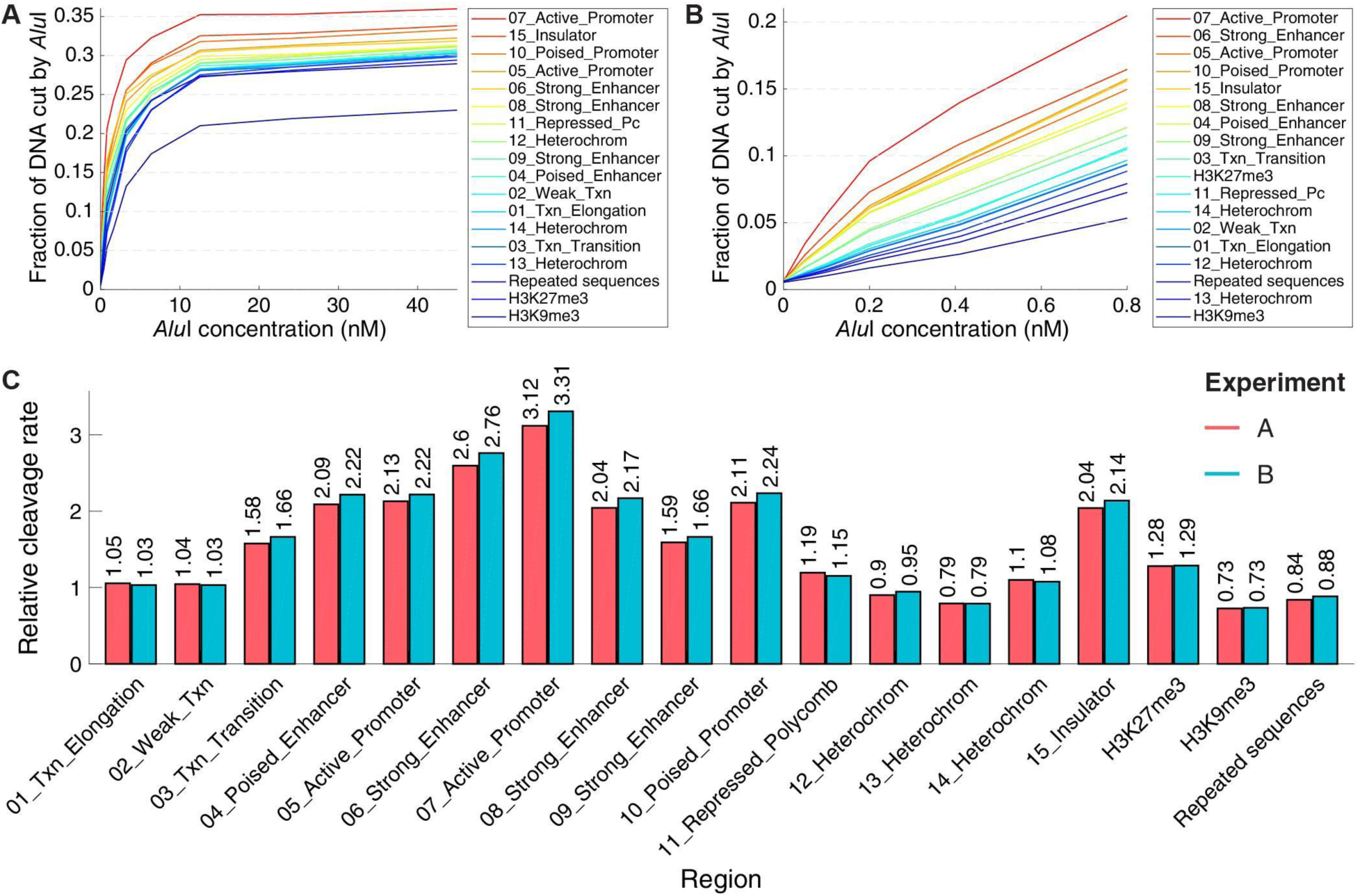
Heterochromatin and euchromatin have very similar DNA accessibilities in mouse hepatocytes. (A) *Alu*I digestion kinetics for sites in the 15 epigenetic chromatin states in mouse hepatocytes defined by Bogu et al. (2015), for sites in repeated sequences (defined by Repeat Masker), and for sites marked by H3K9me3 (constitutive heterochromatin) or H3K27me3 (facultative heterochromatin) (data from Nocetti et al., 2019). (B) Initial rates of *Alu*I digestion. (C) Quantitative comparison of initial digestion rates for the 15 epigenetic states defined by Bogu et al. (2015), repeated sequences and regions marked by H3K9me3 or H3K27me3. Data for biological replicate experiments A and B are shown. See Supplemental Text for details of the analysis.

The hidden Markov model used by Bogu et al. (2015) to define the various chromatin states did not include the histone marks typical of constitutive heterochromatin (H3-K9me2 and H3-K9me3). We confirmed that heterochromatin has a similar accessibility to euchromatin using two independent additional analyses. Firstly, we analyzed H3-K9me3 ChIP-seq data for adult mouse hepatocytes (Nicetto et al. 2019). These data indicate that constitutive heterochromatin defined by the H3-K9me3 mark has a somewhat lower but still quite similar average absolute accessibility (plateau at ∼23%) than the heterochromatin states defined by Bogu et al. (2015) (∼29%; Fig. 5A). In the case of Polycomb-repressed/ facultative heterochromatin, analysis of ChIP-seq data for H3-K27me3 from the same source (Nicetto et al. 2019) indicates a very similar average absolute accessibility (∼28%) to that obtained for the same mark in the Bogu model (state 11; ∼29%; Fig. 5A). Secondly, we used mouse genome annotations to determine the absolute *Alu*I accessibilities of different annotated regions (Supplemental Fig. S4). We observed that all annotated regions have similar accessibilities except promoters (because of their NDRs, as discussed above). Most importantly, repeated sequences, which are strongly enriched in constitutive heterochromatin, have similar average absolute accessibility (∼29%) to that of the heterochromatin states defined by Bogu et al. (∼29%; Fig. 5A).

Although the absolute *Alu*I accessibilities (plateau values) are similar for euchromatin and heterochromatin, it seemed possible that they might be digested at very different rates, reflecting their very different degrees of compaction. Accordingly, we analyzed the initial *Alu*I digestion rates for accessible sites in mouse chromatin (Fig. 5B, C). Regulatory elements containing NDRs (active promoters, insulators and strong enhancers) are digested about 3 times faster than the other chromatin states (*cf*. yeast NDRs; Supplemental Fig. S1). However, accessible sites in heterochromatin (states 11 - 14) are digested at virtually the same rate as those in euchromatin (active genes; states 1 and 2) (Fig. 5B, C). Similarly, the relative rate of *Alu*I digestion of facultative heterochromatin as defined by Bogu et al. (2015) (state 11; ∼1.2-fold) is very similar to that defined by H3-K27me3 ChIP-seq data (∼1.3-fold) (Nicetto et al. 2019). The rate of digestion of constitutive heterochromatin defined by the H3K9me3 mark (∼0.7-fold) is a little slower than for the various heterochromatin states of Bogu et al. (states 12, 13 and 14, which range from ∼0.8 to ∼1.1-fold), but this is a very small effect. If heterochromatin really blocks accessibility, a very large difference in *Alu*I digestion rates for euchromatin and heterochromatin is expected, but it is not observed. Thus, DNA accessibility in mouse hepatocytes does not depend strongly on epigenetic state.

## DISCUSSION

### Nucleosome spacing, phasing and DNA accessibility in chromatin

Our data indicate that access to most of the genome is blocked by nucleosomes in yeast (∼78%) and in mouse hepatocytes (∼68%), consistent with average nucleosome spacings of ∼165 bp and ∼195 bp, respectively, and a protected inner core of ∼133 bp. Thus, nucleosome spacing is the major determinant of absolute DNA accessibility. The inner nucleosome core completely blocks *Alu*I but, since nucleosomes can occupy different positions in different cells (Shen et al. 2001; Cole et al. 2011; Small et al. 2014), all *Alu*I sites are accessible in some cells. Thus, DNA accessibility varies from cell to cell. In yeast, most nucleosomes are phased, but nucleosome positioning is not strong enough to guarantee the inaccessibility of specific sites (Fig. 3). In mouse cells, most nucleosomes are regularly spaced, but are only well-positioned (phased) in the vicinity of active regulatory elements. Although inactive promoters have no NDR or phased nucleosomes, the nucleosomes are still regularly spaced, such that the probability of an *Alu*I site being in the linker is determined by the average spacing.

The plateau in *Alu*I digestion indicates that protection is stable during the 20 minute digestion period, which is inconsistent with widespread nucleosome mobility in isolated nuclei, which would predict continued digestion if nucleosomes slide back and forth, alternately exposing and burying *Alu*I sites. Nucleosomes may be more mobile in vivo due to the activities of various ATP-dependent chromatin remodelers capable of moving nucleosomes. It is likely that isolation of nuclei “freezes” the chromatin structure in the absence of ATP. Nucleosome mobility in vivo would be expected to increase the accessibility of DNA in chromatin.

### DNase I hypersensitivity, ATAC-seq and *Alu*I digestion at promoters

The hypersensitivity of active promoters to DNase I and the contrast between active and inactive promoters (compare the top and bottom halves of the heat map in Fig. 4C) suggests a very large difference in promoter accessibility between active and inactive promoters. In contrast, our *Alu*I data indicate that the difference in absolute accessibility is quite small: ∼58% at the average active promoter NDR compared with ∼32% at inactive promoters (which have no NDR) (Fig. 4D). Similarly, our data also reveal that the difference in initial *Alu*I digestion rates between active promoters and chromatin lacking NDRs is only ∼3-fold (Fig. 5B, C). To reconcile these apparently different results, we note that DNase I hypersensitivity correlates with the presence of a promoter NDR (MNase-seq data; Fig. 4C) and that DNase I data derive from short DNA fragments released at a very early stage in digestion and so are heavily enriched for open chromatin states (NDRs); the rest of the genome is not sequenced. DNA fragments from open promoters are therefore amplified relative to the rest of the genome, resulting in a large artificial difference in accessibility between active and inactive promoters. Similar considerations apply to all methods that sequence only the initial digestion products, including typical ATAC-seq experiments. In our qDA-seq method, all of the DNA fragments are sequenced. We also perform an enzyme titration to prove that the accessibility limit (plateau) has been reached, but this is not possible with DNase-seq, since DNase I is not completely blocked by nucleosomes.

More generally, the DNase I hypersensitivity and Tn5 transposase (ATAC-seq) sensitivity of a promoter depend on two factors: (i) The fraction of accessible promoters: An *Alu*I site in a specific active, open promoter is accessible in some cells but not in the others (information not provided by DNase I or ATAC-seq data). The higher this fraction is, the more DNase I or transposase cutting there would be. (ii) The width of the NDR (target size): The wider the NDR, the higher the probability of DNase I or transposase cleavage, because they are non-specific nucleases (sequence preference may be another important factor). This is clear from the heat maps in Fig. 4, in which the promoters with the most DNase I cleavage are also the ones with the widest NDRs (compare the tops of the DNase I and MNase-seq heat maps). A promoter is unlikely to contain more than one *Alu*I site and so the target size is a much less important factor.

Alternative methods for quantitative measurement of DNA accessibility involve the use of DNA methyltransferases. These enzymes can be used to methylate cytosines in accessible CpG or GpC dinucleotides, which can then be identified by their resistance to bisulphite conversion after sequencing (e.g. NoME-seq (Kelly et al. 2012) or MAPit (Nabilsi et al. 2014)). The methylation pattern reveals the footprints of nucleosomes and other stably bound proteins and therefore has much higher resolution than qDA-seq, although it is not as good as MNase-seq, and may be confounded by natural CpG methylation in the cells of higher organisms. This approach requires considerably more sequence coverage than qDA-seq and the bioinformatic analysis is much more complex. As for qDA-seq, it is necessary to demonstrate that a plateau has been reached before the inaccessible fraction can be inferred.

### Restriction enzymes as proxies for sequence-specific transcription factors

Since restriction enzymes are sequence-specific, they may be considered proxies for transcription factors. Sequence-specific transcription factors must search the DNA sequence to find their cognate binding sites. The search process is facilitated by one-dimensional diffusion of the transcription factor along the DNA in non-specific binding mode, with occasional dissociation and re-association events (Halford and Marko 2004; Woringer and Darzacq 2018). Both transcription factors and restriction enzymes find their cognate sites using this type of mechanism. When a transcription factor locates a cognate site, it remains bound for a relatively long time and may recruit other factors. A restriction enzyme locates a cognate site in the same way, but then cuts the DNA instead, providing a record of that binding event, which we detect and quantify in our experiment. We note that the *Alu*I concentration range used in our experiments is in the expected range for transcription factors (up to ∼50 nM).

### Mouse promoter accessibility and gene repression

The accessibility of *Alu*I sites in inactive hepatocyte promoters implies that transcription factors can bind their cognate sites in inactive promoters in some cells, depending on whether the site is in a linker or not, and that gene inactivity is not primarily due to binding site occlusion. Although cognate sites in inactive promoters are accessible in some cells because they are located in linker DNA, an NDR is not created, suggesting that transcription factor access to DNA is insufficient for gene activation. Pioneer factors (defined as sequence-specific transcription factors capable of binding nucleosomal sites with high affinity (Zaret and Mango 2016)) may be critical, because they have the potential to bind their sites in all cells in a population, whether or not they are occupied by a nucleosome. If so, gene activation would depend on whether the pioneer factor is expressed. However, for reasons that are unclear, some pioneer factors cannot access all of their sites in vivo (Donaghey et al. 2018). An alternative model is that the key to NDR formation may be the clustering of transcription factor binding sites at promoters and enhancers; several specific transcription factors may have to be expressed and bind in concert to form an NDR before a gene can be activated. In this cooperative multi-site model (Adams and Workman 1995), single factor binding events at cognate sites in linker DNA would not be sufficient for NDR formation; all of the factors involved would need to be expressed to initiate activation.

### Heterochromatin and euchromatin have similar DNA accessibilities

Our data indicate that heterochromatin is not generally less accessible than euchromatin. This conclusion is consistent with a recent quantitative analysis of MNase-seq data for human cells (Schwartz et al. 2018). We note that some accessibility is expected given that constitutive heterochromatin must be transcribed to produce the RNA required for its repression (Grewal 2010). Our data indicate that transcription factors would be expected to penetrate heterochromatin, even in its extremely compact state. However, the size of the transcription factor may be critical. Although *Alu*I, which is a monomer with a relative molecular mass of ∼46,000, is similar in size to many transcription factors, theoretical modeling suggests that much larger complexes may be excluded from compact heterochromatin (Maeshima et al. 2015). This is a distinct possibility given that many transcription factors are associated with large complexes.

## METHODS

### *Alu*I digestion of yeast nuclei

Yeast strain YDC111 (***MATa*** *ade2-1 can1-100 leu2-3,112 trp1-1 ura3-1* (Kim et al. 2006)) was grown at 30°C in synthetic complete (SC) medium to A_600_ = ∼0.2 and arrested in G1 by addition of α-factor (FDA Core Facility) to 10 μg/ml. Arrest was monitored by observing the appearance of the “shmoo” phenotype in a light microscope. After 2 h, the cells were harvested by filtration and stored at −80°C. Spheroplasts were prepared from ∼100 A_600_ units of cells in 15 ml SM Buffer (SC medium with 1 M sorbitol, 50 mM Tris-HCl pH 8.0, 20 mM 2-mercaptoethanol) by digestion with ∼26,000 units of lyticase (Sigma L-2524) at 30°C for 5 min or less. Digestion of the cell wall was monitored by measuring the A_600_ of 30 μl cell suspension in 1 ml 1% SDS and considered complete when the A_600_ decreased to < 10% of the initial value. For 3AT experiments, YDC111 cells were grown to mid-log phase at 30°C either in SC medium lacking histidine, followed by addition of 3AT (Sigma 61-82-5) to 10 mM for 20 min, or in SC medium (control), and stored as above. Spheroplasting of 3AT-treated cells was carried out in SM medium lacking histidine. Spheroplasts were spun down in a pre-cooled Sorvall SA600 rotor (7,500 rpm for 5 min at 4°C) and washed once with 25 ml cold ST Buffer (1 M sorbitol, 50 mM Tris-HCl pH 8.0). Spheroplasts were lysed by resuspension in 20 ml cold F Buffer (18% w/v Ficoll-PM400 (GE Healthcare 17-0300-50), 40 mM potassium phosphate, 1 mM MgCl_2_, pH 6.5; protease inhibitors (Roche 05056489001) and 5 mM 2-mercaptoethanol were added just before use). The lysate was applied to a step gradient of 15 ml cold FG Buffer (7% w/v Ficoll-PM400, 20% glycerol, 40 mM potassium phosphate, 1 mM MgCl_2_, pH 6.5, with protease inhibitors and 5 mM 2-mercaptoethanol as above) and spun in an SA600 rotor (12,500 rpm for 20 min at 4°C). The pellets (crude nuclei) were resuspended in 4.4 ml pre-warmed *Alu*I Digestion Buffer (10 mM HEPES pH 7.5, 35 mM NaCl, 5 mM MgCl_2_, with protease inhibitors and 5 mM 2-mercaptoethanol) and divided into eleven 400 μl aliquots. *Alu*I (New England Biolabs R0137 at 0.015 mg/ml; MW = 46,000) was added (0, 1, 2.5, 5, 10, 20, 40, 100, 200, 400, 800 units), mixed thoroughly, and incubated at 25°C for 20 min. Digestion was stopped by adding 50 μl 90 mM EDTA, 9% SDS. Aliquots (180 μl) were removed from each digest to ascertain the level of digestion; the remainders were stored at −20°C prior to sonication. For gel analysis, the DNA was purified by addition of 10 μl 20% SDS, mixing, addition of 50 μl 5 M potassium acetate, followed by two extractions with an equal volume of chloroform, precipitation with 0.7 vol. isopropanol, and one wash with 75% ethanol. The purified DNA was dissolved in 20 μl 10 mM Tris-HCl pH 8.0, 0.1 mM EDTA, 0.5 mg/ml RNase A and incubated at 37°C for 1 h. The DNA was analyzed in a 1% agarose gel stained with SYBR-Gold (Invitrogen S11494). For sonication, the samples were adjusted to 450 μl with 180 μl *Alu*I Digestion Buffer, transferred to 15-ml Sumilon TPX tubes (Diagenode C30010009) and sonicated using a Diagenode Bioruptor 300 at 4°C and high power: 20 cycles of 30 s on and 30 s off. The DNA was purified, treated with RNase as above, purified again using Qiagen PCR purification columns (Qiagen 28106) and eluted in 50 μl 10 mM Tris-HCl pH 8.0, 0.1 mM EDTA (TE(0.1)). Concentrations were determined by measuring A_260_. The degree of sonication was checked by analysis in a 2% agarose gel stained with SYBR-Gold; DNA sizes ranged from ∼100 to ∼700 bp. Prior to library preparation, the DNA was treated with repair enzymes (New England Biolabs PreCR kit M0309) and purified using Qiagen PCR purification columns as above.

### *Alu*I digestion of mouse liver nuclei

Livers were dissected from pregnant (E13.5) female NIH/Swiss mice and stored at −80°C. For each experiment, a liver was thawed on ice and gently disrupted in a glass homogeniser containing 12 ml cold Buffer A per gram liver (Buffer A: 0.34 M sucrose, 60 mM KCl, 15 mM NaCl, 15 mM Tris-HCl pH 8.0, 0.5 mM spermidine-HCl, 0.15 mM spermine, 1 mM Na-EDTA, 15 mM 2-mercaptoethanol, and protease inhibitors as above). The homogenate was filtered through four layers of cheesecloth into a 50-ml tube on ice. Crude nuclei were collected by applying the filtrate to two 4-ml step gradients of Buffer A with 1 M sucrose in 15-ml tubes and spinning in a Sorvall SA600 rotor at 12,500 rpm for 15 mins at 4°C. The supernatants were decanted and solid material on the tube sides was removed with a tissue. The nuclei were washed by gentle resuspension of both pellets in a total of 5 ml Buffer A and spun for 5 min as above. The supernatant was removed, the pellet resuspended in 1 ml Buffer A, and placed on ice. The DNA concentration was estimated by measuring the A_260_ of 2 μl nuclei in 1 ml 1 M NaOH. The volume of nuclei corresponding to 50 A_260_ units was transferred to a 1.5-ml microfuge tube and spun for 5 min at 14,000 rpm at 4°C. The supernatant was removed and the nuclei were resuspended in 1 ml Mouse *Alu*I Digestion Buffer (0.34 M sucrose, 60 mM KCl, 15 mM NaCl, 15 mM Tris-HCl pH 8.0, 5 mM MgCl_2_, 15 mM 2-mercaptoethanol, with protease inhibitors as above). The A_260_ of the diluted nuclei (10 μl) was measured as above. Twelve aliquots of carefully resuspended nuclei, each containing 20 μg DNA in 200 μl Digestion Buffer, were titrated with *Alu*I as follows: 0, 0.3, 0.6, 1.3, 2.5, 5, 10, 20, 40, 80, 160, 320 units. The samples were mixed gently but thoroughly with a 1-ml pipette and incubated at 37°C for 20 min. Digestion was stopped by adding 200 μl 2% SDS, 20 mM EDTA, 10 mM Tris-HCl pH 8.0, mixing thoroughly and incubating for 40 min at room temperature to ensure complete protein removal from DNA. The extent of digestion was determined by analysis of DNA purified from 50 μl of each digest in an agarose gel, after RNase treatment as above. The remaining 350 μl was stored at −20°C prior to sonication. The samples were warmed to room temperature to dissolve precipitated SDS, the volumes were adjusted to 450 μl with 100 μl 10 mM TE(0.1) and sonicated as above. The DNA was purified as above and dissolved in 45 μl 50 mM Tris-HCl pH 8.0, 5 mM EDTA, 0.1 mg/ml RNase A and incubated for 1 h at 37°C. The salt concentration was adjusted by addition of 5 μl 10x NEB Buffer 4 and the DNA was purified using Qiagen PCR columns. DNA was eluted in 50 μl TE(0.1). Concentrations were measured by A_260_. The degree of sonication was checked by analysis in a 2% agarose gel; DNA sizes ranged from ∼100 to ∼700 bp.

### Illumina paired-end library preparation

The Illumina paired-end adaptor was ligated to ∼500 ng purified sonicated *Alu*I-digested DNA using the NEBNext Ultra DNA library kit for Illumina (New England Biolabs E7370) according to the manufacturer’s instructions. The ligated DNA samples were purified without size selection using AMPure XP beads (Beckman A63880). The DNA (50-100 ng) was amplified using the Phusion Hi-Fi PCR master mix with HF buffer (New England Biolabs M0531) or the Q5 Hot Start HiFi PCR master mix (New England Biolabs E6625AA) (7 - 10 cycles). Library quality was checked in an agarose gel. Sequencing was performed using either an Illumina HiSeq-2500 or an Illumina NextSeq-500.

### Bioinformatics and data analysis

Paired-end reads were aligned to the *S. cerevisiae* reference genome sacCer3, or to the *M. musculus* reference genome mm10, using Bowtie2 (Langmead and Salzberg 2012) with the parameters -X 5000 --very-sensitive, to map sequences up to 5 kb with maximum accuracy. For every *Alu*I motif (AGCT) found in the genome, we estimated the fraction of nuclei in which the given motif was cleaved by *Alu*I, *f*_*cut*_ *= N*_*cut*_*/(N*_*cut*_ *+ N*_*uncut*_), by counting the number of reads that were cut at this site (with an end at this site), *N*_*cut*_, and the number of reads that were not cut at this site (overlapping the motif), *N*_*uncut*_, using the Bioinformatics toolbox from Matlab. Because we sequenced 50 nucleotides from both ends of the DNA fragments, we discarded the *Alu*I sites which were less than 50 bp apart, as the reads originating from cleavages at both sites were underrepresented in the properly aligned reads. GEO database data used: MNase-Exo-seq (GSE65889), DNase-seq (GSM1014183 in GSE37074) and RNA-seq (GSM2071423 and GSM2071424 in GSE78391).

## DATA ACCESS

The sequencing data from this study have been submitted to the NCBI Gene Expression Omnibus (http://www.ncbi.nlm.nih.gov/geo/) under accession number GSE115693. Reviewer link: https://www.ncbi.nlm.nih.gov/geo/query/acc.cgi?token=crszoicibhuxvkd&acc=GSE115693 Code: https://github.com/rchereji/AluI_accessibility_analysis (code from this repository will be made public before publication).

### ACKNOWLEDGMENTS

We thank Gary Felsenfeld, Gordon Hager, Alan Hinnebusch, Tom Johnson, and Diego Presman for comments on the manuscript, Valya Russanova for technical support, and Phil Lee, Will Huffman and Doug Fields for mouse livers. We thank the NHLBI Sequencing Core Facility (Yan Luo, Poching Liu and Yuesheng Li) for paired-end sequencing. This study utilized the computational resources of the NIH HPC Biowulf cluster. This work was supported by the Intramural Research Program of the National Institutes of Health (NICHD).

