## Supplemental Information for "DNA accessibility is not the primary determinant of chromatin-mediated gene regulation"

**Supplementary References** **15**

### Supplementary Text

#### Restriction enzyme digestion kinetics

The kinetics of enzyme processes can be investigated using the Michaelis–Menten framework, which we briefly describe in this section. Let's consider the following enzyme catalyzed reaction,

$$E + S \underset{k_{-1}}{\overset{k_1}{\rightleftharpoons}} ES \xrightarrow{k_2} E + P,$$

where we denote the enzyme (*Alu*I restriction enzyme in our case) with $E$, the DNA containing the *Alu*I target site ("substrate") with $S$, the enzyme-substrate complex (enzyme bound to its site) by $ES$, and the cut DNA ("product") with $P$. The first step of association between the substrate and enzyme to form the complex $ES$ is reversible, with forward and reverse rates $k_1$ and $k_{-1}$, respectively. The second step of DNA cleavage by *Alu*I is assumed to be irreversible, happening with the rate $k_2$.

Let's denote the concentrations of each component as follows:

$$\begin{aligned}
[E] &- \text{the concentration of free enzyme } E;\\
[S] &- \text{the concentration of substrate } S;\\
[ES] &- \text{the concentration of enzyme-substrate complex } ES;\\
[P] &- \text{the concentration of product } P.
\end{aligned}$$

These concentrations satisfy the following reaction equations:

$$\frac{\mathrm{d}[S]}{\mathrm{d}t} = -k_1[S][E] + k_{-1}[ES], \tag{1}$$

$$\frac{\mathrm{d}[E]}{\mathrm{d}t} = -k_1[S][E] + (k_{-1} + k_2)[ES], \tag{2}$$

$$\frac{\mathrm{d}[ES]}{\mathrm{d}t} = k_1[S][E] - (k_{-1} + k_2)[ES], \tag{3}$$

$$\frac{\mathrm{d}[P]}{\mathrm{d}t} = k_2[ES]. \tag{4}$$

Adding Eqs. (2) and (3) we obtain that

$$\frac{\mathrm{d}[E]}{\mathrm{d}t} + \frac{\mathrm{d}[ES]}{\mathrm{d}t} = 0,$$

meaning that $[E](t) + [ES](t) = [E]_0$, so the total concentration of enzyme (either free or bound to DNA) remains constant during the experiment. Adding Eqs. (1), (3), and (4) we obtain that

$$\frac{\mathrm{d}[S]}{\mathrm{d}t} + \frac{\mathrm{d}[ES]}{\mathrm{d}t} + \frac{\mathrm{d}[P]}{\mathrm{d}t} = 0,$$

so the total amount of DNA (either cut or uncut) is also constant, $[S](t) + [ES](t) + [P](t) = [S]_0$. Therefore, we can solve for $[E](t)$ and $[P](t)$,

$$\begin{aligned}
[E](t) &= [E]_0 - [ES](t),\\
[P](t) &= [S]_0 - [S](t) - [ES](t),
\end{aligned}$$

and substitute these into the initial system of four differential equations, which reduces to:

$$\frac{\mathrm{d}[S]}{\mathrm{d}t} = -k_1[S]([E]_0 - [ES]) + k_{-1}[ES],$$
$$\frac{\mathrm{d}[ES]}{\mathrm{d}t} = k_1[S]([E]_0 - [ES]) - (k_{-1} + k_2)[ES].$$

This set of coupled nonlinear differential equations cannot be solved analytically, and numerical integration is required to obtain an accurate solution. However, the situation simplifies considerably if we assume that the cleavage reaction happens very fast compared to the binding and unbinding processes, i.e. if we assume that the process of searching the specific site of *Alu*I is the rate limiting step. In this case, we can use the quasi-steady-state approximation (QSSA), which assumes that the reactive intermediate $ES$ will be present at a low concentration that is changing very slowly throughout the course of the reaction, i.e. $\frac{\mathrm{d}[ES]}{\mathrm{d}t} \approx 0$. This type of equations and assumptions can be rigorously studied using singular perturbation theory, which is outside the scope of this article.

Using the QSSA we obtain

$$k_1[S]([E]_0 - [ES]) - (k_{-1} + k_2)[ES] = 0,$$

and

$$[ES] = \frac{[E]_0[S]}{K_m + [S]}$$

where $K_m = \frac{k_{-1}+k_2}{k_1}$ is known as the Michaelis constant. From Eq. (4) we obtain the production rate

$$\frac{\mathrm{d}[P]}{\mathrm{d}t} = k_2[ES] = \frac{V_{\max}[S]}{K_m + [S]},$$

where $V_{\max} = k_2[E]_0$. Substituting $[ES]$ into the equation for $[S]$ we obtain

$$\begin{aligned}\frac{\mathrm{d}[S]}{\mathrm{d}t} &= -k_1[S]\left([E]_0 - \frac{[E]_0[S]}{K_m + [S]}\right) + k_{-1}\frac{[E]_0[S]}{K_m + [S]} \\ &= \frac{[E]_0[S]}{K_m + [S]}\left(-k_1(K_m + \cancel{[S]}) + \cancel{k_1[S]} + k_{-1}\right) \\ &= -\frac{V_{\max}[S]}{K_m + [S]}.\end{aligned}$$

So, we obtained the Michaelis-Menten equation, which gives the rate of product creation and substrate decay

$$\frac{\mathrm{d}[P]}{\mathrm{d}t} = -\frac{\mathrm{d}[S]}{\mathrm{d}t} = \frac{V_{\max}[S]}{K_m + [S]}, \tag{5}$$

where the maximum rate is $V_{\max} = k_2[E]_0$, and the Michaelis constant is $K_m = \frac{k_{-1}+k_2}{k_1}$.

The Michaelis-Menten rate equation does not have a definite order with respect to the substrate, as in different regimes of the substrate concentration it can be simplified either to a first-order rate equation,

$$\frac{\mathrm{d}[S]}{\mathrm{d}t} \approx -\frac{V_{\max}}{K_m}[S], \text{if } [S] \ll K_m,$$

or to a zero-order rate equation,

$$\frac{\mathrm{d}[S]}{\mathrm{d}t} \approx -V_{\max}, \text{if } [S] \gg K_m.$$

In both cases the rate equation can be easily integrated to obtain

$$[S](t) = [S]_0 \exp\left(-\frac{V_{\max}}{K_m}t\right) \text{ (first-order rate)},$$

or

$$[S](t) = [S]_0 - V_{\max}t \text{ (zero-order rate)},$$

where $[S](0) = [S]_0$.

In the general case, integrating Eq. (5) we obtain

$$\int \frac{K_m + [S]}{[S]}\mathrm{d}[S] = \int -V_{\max}\mathrm{d}t$$

or

$$K_m \log\left(\frac{[S](t)}{[S]_0}\right) + ([S](t) - [S]_0) + V_{\max}t = 0.$$

This is an exact solution of the Michaelis-Menten equation, but unfortunately it is an implicit nonlinear equation. The explicit closed-form solution of this equation (*1, 2*) is rarely used, as it involves the Lambert $W$ function, the special function that solves the transcendental equation

$$W(x)e^{W(x)} = x.$$

The explicit solution of the Michaelis-Menten rate equation (*1, 2*) is

$$[S](t) = K_m W\left(\frac{[S]_0}{K_m}\exp\left(\frac{[S]_0 - V_{\max}t}{K_m}\right)\right), \tag{6}$$

which can be conveniently rewritten in a dimensionless form as

$$s(t) = W\left(s_0 \exp\left(s_0 - vt\right)\right), \tag{7}$$

where $s = [S]/K_m$ is the reduced concentration, and $v = V_{\max}/K_m$ is the first-order rate constant, which is directly proportional to the enzyme concentration,

$$v = \frac{V_{\max}}{K_m} = \frac{k_1 k_2}{k_{-1} + k_2}[E]_0.$$

In our experiments, instead of digesting the chromatin for different amounts of time, we digested the chromatin for a fixed time, $T = 20$ min, but using different concentrations of *Alu*I, $[E]$. In this case, the reduced concentration of substrate at time $t = T$ can be written as a function of the enzyme concentration that was used in a particular experiment,

$$\begin{aligned} s([E]) &= W\left(s_0 \exp\left(s_0 - v([E])T\right)\right) \\ &= W\left(s_0 \exp\left(s_0 - k[E]\right)\right), \end{aligned} \tag{8}$$

which has a simple form, with only two free parameters:

$$k = \frac{v([E])T}{[E]} = \frac{k_1 k_2}{k_{-1} + k_2} T, \tag{9}$$
$$s_0 = \frac{[S]_0}{K_m}.$$

In a population of cells, a specific *Alu*I site can be either accessible to the restriction enzyme, or inaccessible – blocked by nucleosomes or other DNA-binding proteins. Let $A$ be the fraction of cells in which this *Alu*I site is accessible and $1 - A$ the inaccessible fraction. Let's denote the concentration of accessible and inaccessible *Alu*I sites by

$[D]$ – the concentration of accessible sites (free DNA);

$[B]$ – the concentration of inaccessible/blocked sites.

Then, at the beginning of the reaction we have that

$$[D](0) = [D]_0 = A[S]_0,$$
$$[B](0) = [B]_0 = (1 - A)[S]_0,$$

and the corresponding dimensionless variables

$$d_0 = \frac{[D]_0}{K_m} = As_0,$$
$$b_0 = \frac{[B]_0}{K_m} = (1 - A)s_0.$$

At the end of the digestion, according to Eq. (8) we have that

$$\begin{aligned} d([E]) &= W\left(d_0 \exp\left(d_0 - k[E]\right)\right) \\ &= W\left(As_0 \exp\left(As_0 - k[E]\right)\right), \end{aligned}$$

while the amount of inaccessible/blocked substrate remains constant during the cleavage reaction,

$$b([E]) = b_0 = (1 - A)s_0.$$

The fraction of *Alu*I sites that remain uncut after time $T$ is

$$\begin{aligned} f_{\text{uncut}}([E]) &= \frac{d([E]) + b([E])}{d_0 + b_0} \\ &= \frac{W\left(As_0 \exp\left(As_0 - k[E]\right)\right) + (1 - A)s_0}{s_0} \\ &= \frac{W\left(As_0 \exp\left(As_0 - k[E]\right)\right)}{s_0} + (1 - A). \end{aligned}$$

The fraction of *Alu*I sites that were cut before time $T$ is

$$\begin{aligned} f_{\text{cut}}([E]) &= 1 - f_{\text{uncut}}([E]) \\ &= A - \frac{W\left(As_0 \exp\left(As_0 - k[E]\right)\right)}{s_0}. \end{aligned}$$

So, the fractions of *Alu*I sites that are cut and uncut at the end of the experiment, depend on four variables: $[E]$ – the concentration of restriction enzyme used in the experiment; $A$ – the fraction of accessible sites; $k$ – the first-order cleavage rate; and $s_0$ – the substrate concentration. We obtained that

$$f_{\text{cut}}([E], A, k, s_0) = A\left(1 - \frac{W\left(As_0 \exp\left(As_0 - k[E]\right)\right)}{As_0}\right) \tag{10}$$

$$f_{\text{uncut}}([E], A, k, s_0) = 1 - A + A\frac{W\left(As_0 \exp\left(As_0 - k[E]\right)\right)}{As_0} \tag{11}$$

In Fig. S5 we show the dependence of $f_{\text{cut}}$ and $f_{\text{uncut}}$ on the four parameters: $[E]$, $A$, $k$, and $s_0$. We see that the accessibility parameter, $A$, dictates the asymptotic (plateau) values for $f_{\text{cut}}$ and $f_{\text{uncut}}$ (Fig. S5A):

$$\lim_{[E]\to\infty} f_{\text{cut}} = A,$$

$$\lim_{[E]\to\infty} f_{\text{uncut}} = 1 - A.$$

The rate parameter $k$ dictates the initial slopes of the $f_{\text{cut}}$ and $f_{\text{uncut}}$ curves (Fig. S5B), and the $s_0$ parameter determines the overall shape of the curves (Fig. S5C): a low $s_0$ results in a first-order reaction (exponential decay of the substrate), while a high $s_0$ results in a zero-order reaction (linear decay of the substrate).

Fig. S6 shows the fraction of *Alu*I sites that were (Fig. S6A) or were not cut (Fig. S6B) when different concentrations of *Alu*I were used in the cleavage reaction. Although there is a significant variability among the plateau levels for different sites (percentiles represented as different shades of red and blue in Figs. S6A and S6B, respectively), the overall shape of the medians suggest an exponential decay of the concentration of accessible *Alu*I sites, consistent with a first-order decay reaction. By fitting the medians of $f_{\text{cut}}$ and $f_{\text{uncut}}$ using Eqs. (10) and (11), respectively, we obtained the following fitted parameters: $A = 0.209$, $k = 0.048$, and a very small $s_0$ parameter ($s_0 \ll 1$).

As we pointed out before, when $s_0 \ll 1$ (or equivalently $[S] \ll K_m$), the Michaelis-Menten rate equation becomes a first-order rate equation, with the solution

$$[S](t) = [S]_0 \exp\left(-\frac{V_{\max}}{K_m}t\right).$$

At the end of the reaction (at time $T$), the reduced concentration as a function of the enzyme concentration $[E]$ is

$$s([E]) = s_0 \exp\left(-k[E]\right),$$

where $s = [S]/K_m$ and $k$ is given by Eq. (9). Similarly, in the case of two types of substrate (accessible and blocked sites), we obtain that

$$d([E]) = d_0 \exp\left(-k[E]\right) = As_0 \exp\left(-k[E]\right)$$

and

$$b([E]) = b_0 = (1 - A)s_0.$$

Therefore, the fraction of DNA that was not cut by *Alu*I during the cleavage reaction becomes

$$f_{\text{uncut}}([E], A, k) = \frac{d([E]) + b([E])}{s_0} = Ae^{-k[E]} + (1 - A), \tag{12}$$

while the cut fraction is

$$f_{\text{cut}}([E], A, k) = 1 - f_{\text{uncut}}([E], A, k) = A\left(1 - e^{-k[E]}\right). \tag{13}$$

Clearly, the asymptotic values of these two fractions are

$$\lim_{[E]\to\infty} f_{\text{cut}} = A,$$
$$\lim_{[E]\to\infty} f_{\text{uncut}} = 1 - A,$$

and the accessibility parameter $A$ can be easily obtained from these asymptotes. The other parameter, the first-order rate constant $k$, can be obtained by analyzing the initial slope of $\log(f_{\text{uncut}})$. We have that

$$\log\left(f_{\text{uncut}}([E], A, k)\right) = \log\left(Ae^{-k[E]} + (1 - A)\right).$$

When $[E] \to 0$, we have that

$$e^{-k[E]} \approx 1 - k[E],$$

and

$$\begin{aligned} \log\left(f_{\text{uncut}}([E], A, k)\right) &\approx \log\left(A(1 - k[E]) + (1 - A)\right) \\ &= \log\left(1 - kA[E]\right)) \\ &\approx -kA[E] \end{aligned}$$

so for the initial stages of digestion, $\log(f_{\text{uncut}})$ decreases linearly with $[E]$, with a slope given by

$$\text{Slope} = \frac{\partial}{\partial[E]} \log(f_{\text{uncut}}) = -kA \tag{14}$$

Using the plateau level for $f_{\text{cut}}$ or $f_{\text{uncut}}$ we obtain a quantitative estimation for DNA accessibility $A$, which is not available through other methods, such as MNase-seq, ATAC-seq, or DNase-seq. Knowing the accessibility of each *Alu*I site and measuring the initial slope of $\log(f_{\text{uncut}})$, using Eq. (14) we can compute the rate constant $k$ for each site. If different *Alu*I sites are located in genomic regions characterized by different chromatin openness, then we expect the overall rates $k$ to vary from region to region: *Alu*I sites located in open chromatin will be characterized by higher cleavage rates $k$, while the sites located in closed chromatin will be characterized by lower rates $k$. Surprisingly, when different mouse liver chromatin states were analyzed to determine their typical digestion rate constants, we observed only a relatively small difference among the respective $k$'s (less than 5-fold difference) and not the expected orders-of-magnitude difference. This indicates that the accessibilities of euchromatin and heterochromatin are very similar. Therefore, DNA accessibility is *not* the primary determinant of chromatin-mediated gene regulation.

#### Supplementary Figures

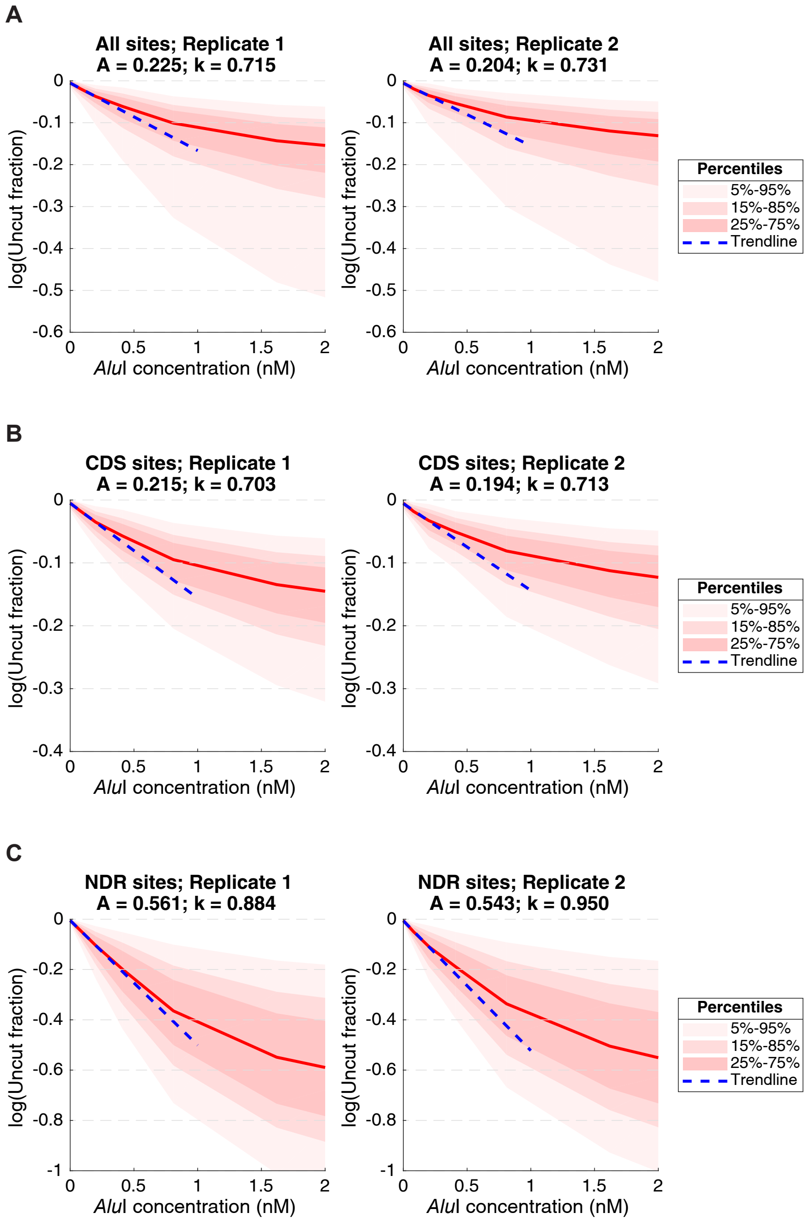

**Fig. S1. Mean initial digestion rates of *Alu*I sites in yeast cells.** The natural logarithm of the uncut fraction vs. *Alu*I concentration for two biological replicate experiments. **(A)** All *Alu*I sites. **(B)** *Alu*I sites in gene bodies. **(C)** *Alu*I sites in promoter NDRs (defined by the locations of the +1 and -1 nucleosomes obtained from (*3*)). See Supplementary Text for a full mathematical treatment.

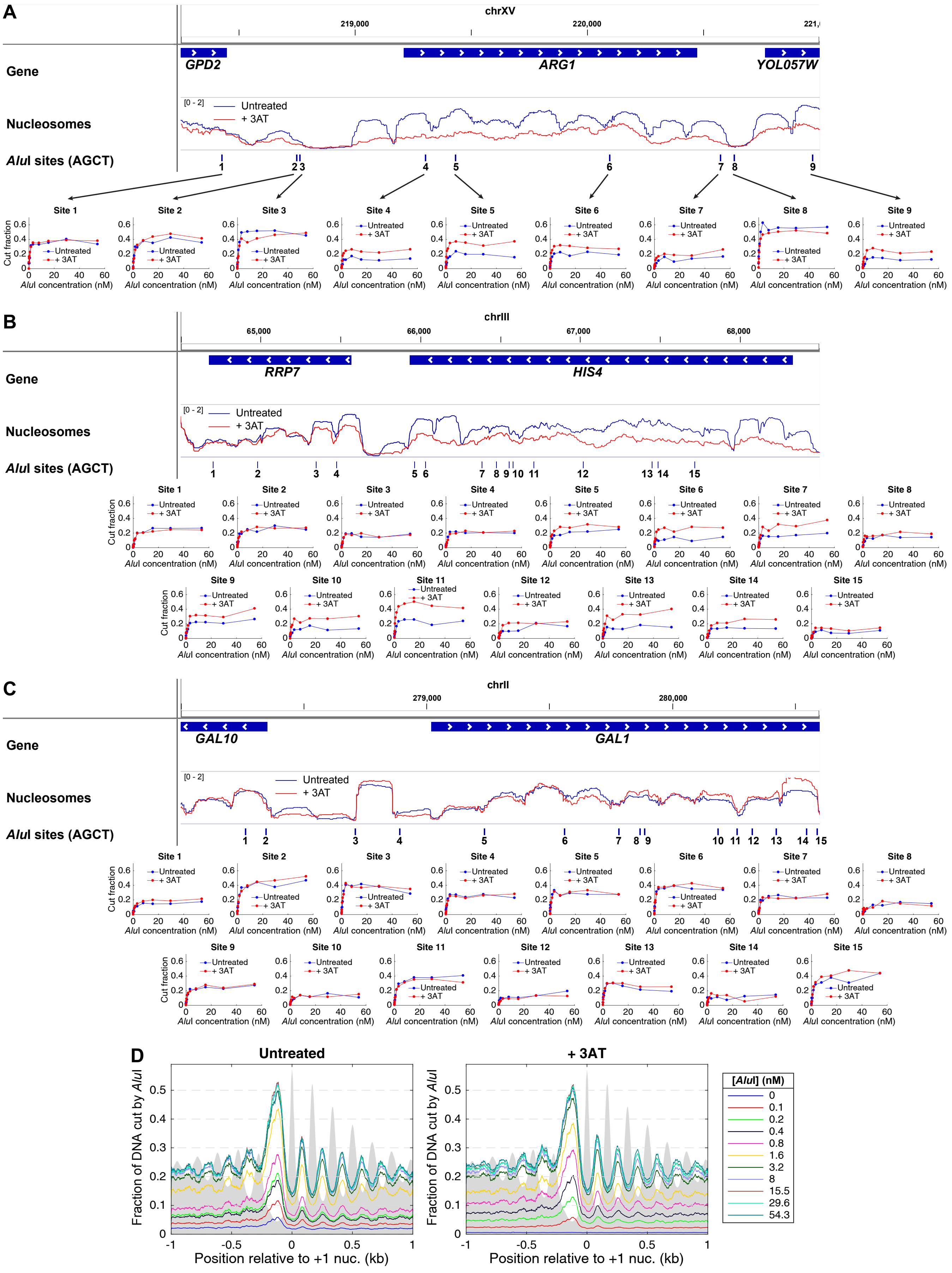

**Fig. S2.** Caption on the next page.

**Fig. S2. Heavy transcription after 3AT induction results in increased *Alu*I accessibility in yeast.** After induction with 3AT, a small number of Gcn4-dependent genes, including **(A)** *ARG1* and **(B)** *HIS4*, show significant nucleosome loss over the coding region, sometimes extending into the flanking regions, as well as disrupted nucleosome spacing and a wider NDR, all of which correlate with heavy transcription (*4,5*). *Alu*I accessibility data and MNase-seq data are shown (blue line: control; red line: +3AT). **(C)** *GAL1* (not induced by 3AT). **(D)** Mean *Alu*I accessibility as a function of distance from the center of the +1 nucleosome (defined in (*3*)) on all ~5,000 yeast genes for exponentially growing cells before and after treatment with 3AT for 20 min (cf. arrested cells; Fig. 2C). Grey area: MNase-seq data showing nucleosome dyad distributions (fragment centers) on an arbitrary scale.

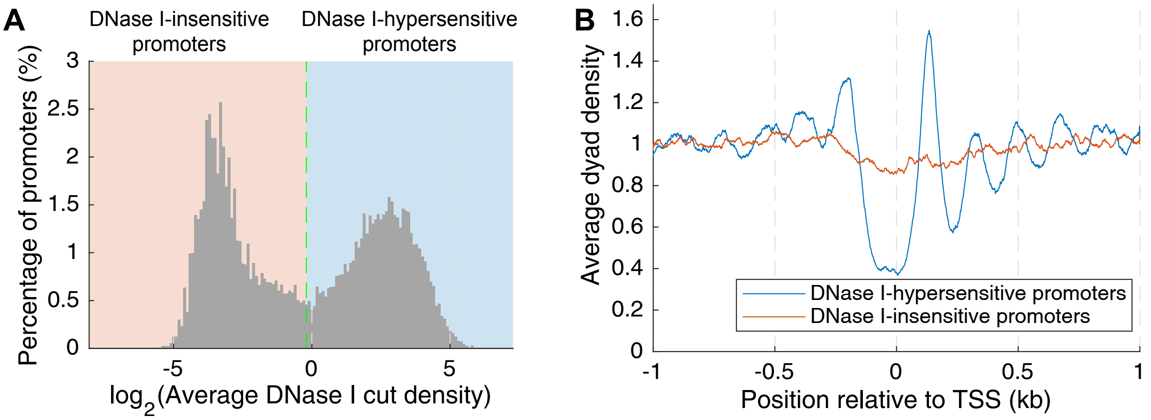

**Fig. S3. Mouse genes in liver can be divided into two distinct chromatin states using promoter DNase I hypersensitivity data. (A)** Histogram of the number of promoters with a given DNase I cut density. The dashed line indicates the dividing line used in the DNase I heatmap in Fig. 3C (white line). **(B)** Nucleosome phasing on genes with DNase I-hypersensitive promoters and DNase I-insensitive promoters (data above and below the white line in the MNase-Exo-seq heatmap in Fig. 3C).

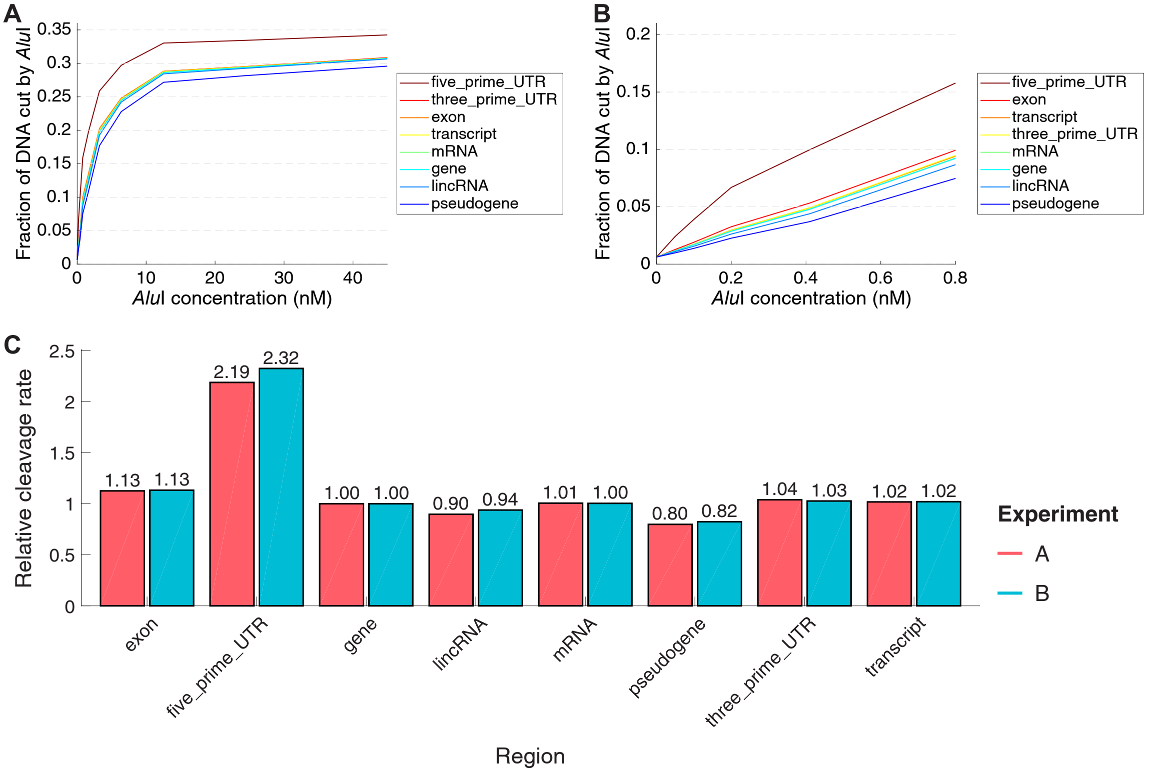

**Fig. S4. Relative *Alu*I cleavage rates for sites in various annotated regions of the mouse genome (mm10).** (A) *Alu*I digestion kinetics for 8 types of annotated regions of the mouse genome. The asymptotic value of $f_{\mathrm{cut}}$ represents the accessible fraction of each region, $A$. (B) Close up view on the domain of low *Alu*I concentrations. The initial slope of each of these curves is equal with $k \cdot A$, so it can be used as a measure of the cleavage rate constant corresponding to each region, $k$. (C) Comparison of the cleavage rate constants corresponding to 8 annotated regions (two biological replicate experiments). See Supplementary Text for details of the analysis.

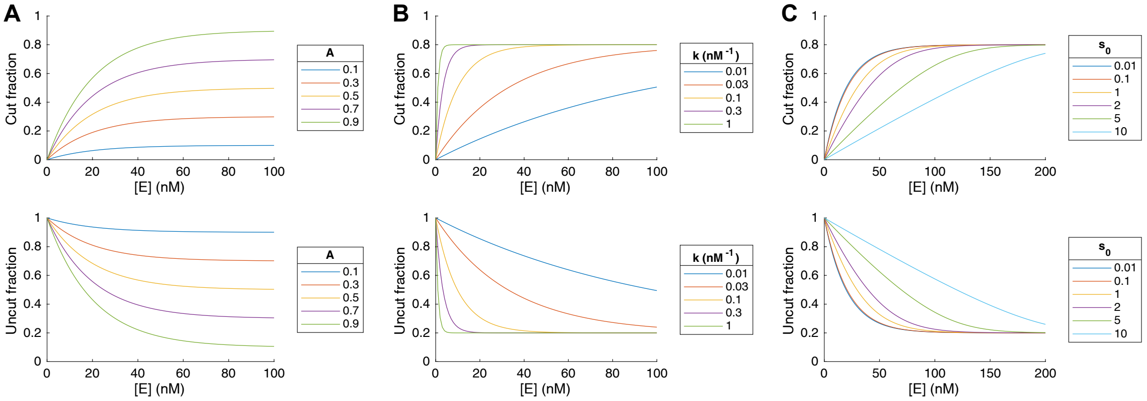

**Fig. S5. Predictions of the exact solution of the Michaelis-Menten equation (Eqs.** (10) **and** (11)**).** The predicted dependence of the fraction of *Alu*I sites that are cut (upper panels) or remain uncut (lower panels) on different parameters: (A) DNA accessibility $A$, (B) reaction rate $k$, and (C) the initial substrate concentration $s_0$ ($[S]_0/K_m$).

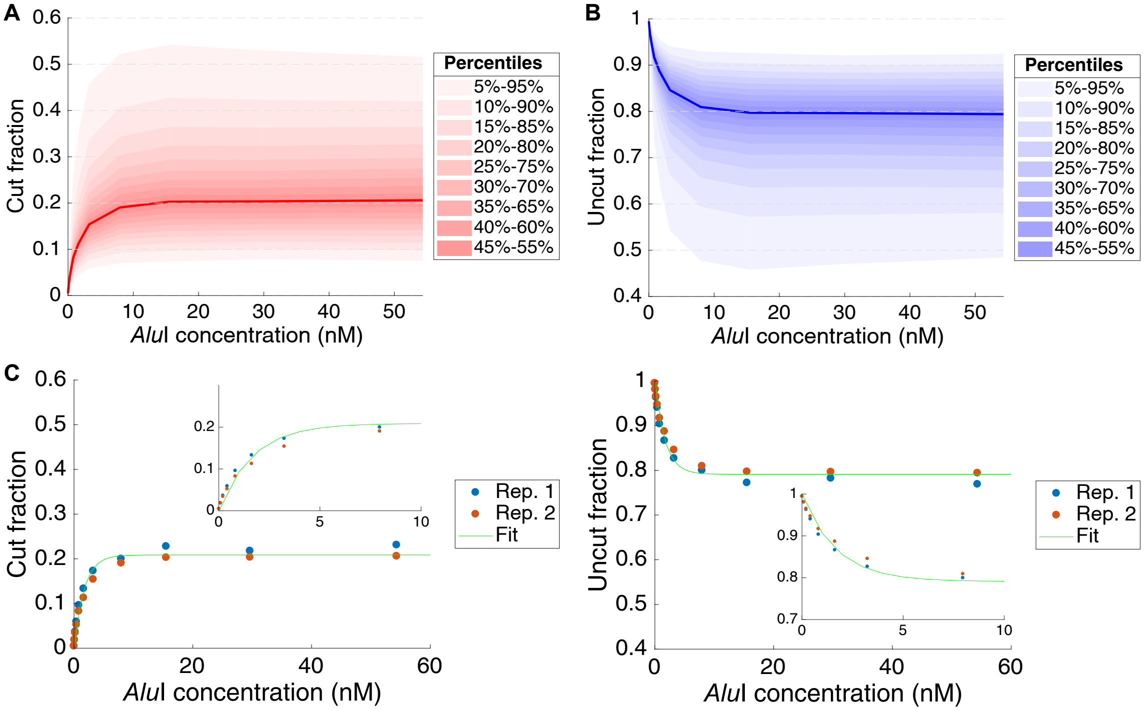

**Fig. S6. Restriction enzyme cleavage in yeast.** (A) The fraction of sites cut by different concentrations of *Alu*I in 20 min. The median cut fraction for each *Alu*I concentration is shown with a red line, and different percentiles of the cut fraction distribution are shown with different shades of red. (B) The fraction of sites that were not cut by *Alu*I after 20 min. The median value is shown with a blue line, and different percentiles are shown with different shades of blue. (C) Fits for the cut and uncut fractions, using the exact solution of the Michaelis-Menten model – Eqs. (10) and (11). Fitted parameters: $A = 0.209$, $k = 0.594$, $s_0 \ll 1$ (too small to get a precise estimate).
